# SARS-COV-2 Recombinant Receptor-Binding-Domain (RBD) Induces Neutralising Antibodies Against Variant Strains of SARS-CoV-2 and SARS-CoV-1

**DOI:** 10.1101/2021.05.10.443438

**Authors:** Lok Man John Law, Michael Logan, Michael Joyce, Abdolamir Landi, Darren Hockman, Kevin Crawford, Janelle Johnson, Gerald LaChance, Holly Saffran, Justin Shields, Eve Hobart, Raelynn Brassard, Elena Arutyunova, Kanti Pabbaraju, Matthew Croxen, Graham Tipples, M Joanne Lemieux, Lorne Tyrrell, Michael Houghton

## Abstract

SARS-CoV-2 is the etiological agent of COVID19. There are currently several licensed vaccines approved for human use and most of them are targeting the spike protein (or virion) in the virion envelope to induce protective immunity. Recently, variants that spread more quickly have emerged. There is evidence that some of these variants are less sensitive to neutralization *in vitro*, but it is not clear whether they can evade vaccine induced protection. In this study, we tested the utility of SARS-CoV-2 spike RBD as a vaccine antigen and explore the effect of formulation with Alum/MPLA or AddaS03 adjuvants. Our results indicate RBD induces high titers of neutralizing antibodies and activates strong cellular immune responses. There is also significant cross-neutralisation of variants B1.1.7 and B.1.351 and to a lesser extent, SARS-CoV-1. These results indicate that recombinant RBD can be a viable candidate as a stand-alone vaccine or as a booster shot to diversify our strategy for COVID19 protection.

## Introduction

Coronaviruses are enveloped positive strand RNA viruses, which are divided into α, β, γ and δ genera, and infect a multitude of host organisms ^1^. Four betacoronaviruses (HCoV-OC43, -HKU1, -NL63, -229E) are endemic in humans and have low pathogenicity. However, three zoonotic coronavirus pathogens (SARS-CoV-1, MERS-CoV and SARS-CoV-2) have emerged into human populations causing fatalities. SARS-CoV-2 shares 79% sequence identify with the original SARS-CoV-1 identified in 2003 ^2^ and is the etiological agent of COVID19. December 2020, approximately a year after the first case was reported, marked the emergency approval of a mRNA based vaccine. It is a tremendous scientific achievement, but challenges remain around the rise of variant strains of SARS-CoV-2. There are examples where prior infection of earlier strain of SARS-CoV-2 cannot prevent infection of new variants ^3,4^. Currently, it is not clear whether the current vaccine can provide sufficient protection to reverse this pandemic. However, there is evidence that declining new infection rates in many places coincide with the introduction of vaccination programs ^3 4,5^. However, *in vitro* testing in the lab, indicates that sera from vaccinated patients exhibits reduced neutralization activity against variants, particularly the variant B.1.351 originally found in South Africa^4–6^. It is not clear how this reduction *in vitro* translates to real-life efficacy.

SARS-CoV-2 spike (S) is the major target for vaccine development^7^. It forms a trimer decorating the surface of virions and is essential for initiating infection by interacting with host receptor angiotensin converting enzyme 2 (ACE-2) followed by membrane fusion ^2,8^. On the virion, the S protein adopts a partial open conformation ^9,10^ ; The opening of S is necessary for efficient interaction with ACE-2 followed by conformational changes that trigger fusion. The spike protein is separated into two functional units. The amino terminal S1 is responsible for binding to host cell receptors and the carboxyl terminal S2 is responsible for mediating fusion of the viral envelope with cellular membranes ^11^. To initiate entry, there is an activation cleavage at the S2’ site, upstream of the fusion peptide. For SARS-CoV-2, the proteases cathepsins D/L or transmembrane protease serine protease-2 (TMPRSS-2) are required ^8,12^. Accordingly, pharmacological agents blocking these proteases can prevent infection *in vitro* ^8,12^. In addition, SARS-CoV-2 has a stretch of polybasic residues between S1 and S2, which is cleaved by the host protease furin ^8,13^. This furin cleavage site is absent in SARS-CoV-1 ^14,15^ and mutation to remove this process has reduced fitness of SARS-COV-2 in cell cultures ^16,17^.

Antibodies targeting S can neutralize and confer protection against SARS-CoV-2 infection. Neutralizing antibodies in convalescent patients correlate with their ability to bind the receptor binding domain (RBD) ^18^, although cellular immune responses are also likely to contribute to protection. The majority of cloned SARS-CoV-2 neutralizing antibodies target the RBD in S1 ^19,20^. Additionally, there are other neutralizing epitopes outside of the RBD in the S1 NTD and S2 domains ^21 22^. Immunogenicity of SARS-CoV-2 RBD has been tested using various expression platforms ^10,23–25^. They are capable of inducing neutralizing antibodies. Comparisons between RBD and full length S in mRNA based vaccine showed comparable immunogenicity in the clinic ^26–28^. The RBD structures of SARS-CoV-2 and SARS-CoV-1 are highly homologous, although S1 sequences are generally more diverse. While there are reports of antibody cross-neutralisation of SARS-CoV-1 and SARS-CoV-2, many monoclonal antibodies against SARS-CoV-1 cannot neutralize SARS-CoV-2 suggesting unique epitopes between these viruses ^29,30^.

In this study, we test the utility of SARS-CoV-2 RBD as a vaccine antigen and explore the effect of formulation with Alum/MPLA or AddaS03 adjuvants. Our results are encouraging; adjuvanted RBD induces high titers of neutralizing antibodies and activates strong cellular immune responses. There is also significant cross-neutralization of variants B1.1.7, B.1.351 and P.1 (originating in U.K., South Africa and Brazil). Therefore, our data suggests that adjuvanted recombinant RBD can be a viable candidate as a stand-alone vaccine or as a booster shot to diversify our global strategy to protect from SARS-CoV-2 infection.

## Materials and Methods

### Cell culture and antibodies

CHO cells stably expressing recombinant RBD of SARS-CoV-2 Spike (aa. 319-591) (GenBank accession no. AAP13567.1) were propagated in ProCho5 media (Lonza) containing glutamine, 1% heat-inactivated fetal bovine serum (FBS) (Thermo Fisher Scientific), and 100 U/ml of penicillin and 100 μg/ml of streptomycin (Pen/Strep; Invitrogen, Carlsbad, CA). 293T cells and Vero E6 cells (ATCC CRL-1586) were propagated in Dulbecco’s modified Eagle’s medium (Thermo Fisher Scientific) containing 10% heat-inactivated fetal bovine serum (Omega Scientific, Tarzana, CA), and Pen/Strep (Invitrogen). 293T cells overexpressing ACE-2 (293T ACE-2) were generously provided by Dr. Paul Bieniasz (The Rockefeller University) ^31^ and cultured in 293T cells media supplemented with 5ug/ml blasticidin. Rabbit anti-SARS-CoV-2 Spike (RBD) antibody was commercially sourced (SinoBiological, Cat# 40592-T62).

### Expression and purification of recombinant SARS-CoV-2 RBD

The SARS-CoV-2 RBD (Genbank AAP13567.1; amino acids 319-591), preceded by the signal peptide sequence for tissue plasminogen activator (tPA) and followed by a human rhinovirus 3C (HRV3C) protease cleavable C-terminal human monomeric IgG1 Fc tag (mFc) was inserted into the SpeI/ XhoI site of the pTRIP lentiviral vector bearing an IRES-AcGFP reporter ^32^. Lentiviral particles were generated in 293T cells according to a previous method ^32^ and used to transduce CHOK1 cells. GFP-positive transduced CHOK1 cells expressing RBD-mFc were sorted by flow cytometry using a BD FACSAria III cell sorter (BD Biosciences) and suspension adapted in PROCHO5 medium (Lonza, Walkersville, MD, USA) with 1 % FBS in shaker flasks (Corning, Corning, NY, USA).

Purification of Recombinant RBD (319-591) was performed with modifications based on a previous published method ^33^. mFc-tagged RBD was captured from CHOK1 cell culture supernatants using mab select sure LX affinity resin (Cytiva, Marlborough, MA, USA), washed with phosphate buffered saline (PBS) and the resin digested with His6-GST-HRV3C protease (Thermo Fisher Scientific) 16-18 hours at 4°C. The digested material was applied to Glutathione Sepharose 4B (Cytiva) to remove the protease and the flow through concentrated using a 30,000 molecular weight cut-off centrifugal filter unit (EMD Millipore, Billerica, MA, USA).

### ACE-2 Expression and Purification

Full length human ACE-2-MycDDK in pCMV-6 entry vector (Origene, cat #RC208442) was expressed in Expi293™ cells. 30 mL cell cultures were grown to a density of 4.5 × 10^6^ – 5.5 × 10^6^ cells/mL and then diluted to a final density of 3 × 10^6^ cells/mL for transfection. The cells were transfected with 1.0 μg plasmid DNA/mL of culture and 80 μL ExpiFectamine™ 293 reagent. Transfection was performed as per the protocol described in the Expi293™ Expression System User Guide (ThermoFisher Scientific). Cells were harvested 4 days post-transfection by centrifugation at 500 x g for 20 min at 4 °C. The cell pellet was resuspended in 30 mL of buffer (50mM Tris-HCl, 300mM NaCl, pH 7.5) containing 1 mM PMSF. Resuspended cells were lysed using 4 passages through an Emulsiflex with a maximum pressure of 30 kPSI. Protein was solubilized by incubating lysed cells with 0.1% Triton x-100 on ice with stirring for 30 minutes. Cellular debris was removed by centrifugation at 20000 x g for 30 min at 4 °C. DDM was added to supernatant to a final concentration of 0.05%.

Anti-FLAG resin was pre-washed with TBS plus 0.05% DDM, then incubated with supernatant on a nutator for 1 hour at 4 °C. The resin was applied to a gravity flow column and the column was then washed with 20 mL of TBS with 0.05% DDM. The protein was eluted in 1 mL aliquots with 0.1 M bicine pH 3.5 into 100 μL of 1M Tris pH 8.0 to neutralize the sample. Protein containing fractions were combined and dialyzed in buffer (50 mM Tris-HCl pH 8.0, 150 mM NaCl, 5% Glycerol) for 6 hours at 4 °C. Concentrated ACE-2-MycDDK was aliquoted, flash frozen with liquid nitrogen and stored at −80 °C for subsequent use.

### Immunization of mice and serum samples

Female CB6F1 mice (Charles River Laboratories, Montreal, QC, Canada) (5-7 weeks old) for vaccination experiments were cared for in accordance with the Canadian Council on Animal Care guidelines. Experimental methods were reviewed and approved by the University of Alberta Health Sciences Animal Welfare Committee. Recombinant RBD (319-591) (1 μg) was mixed either with PBS (Control group), in a 1:1 ratio with 75 μg alum and 7.5 μg monophosphoryl Lipid A (Alum/MPLA group) (Invivogen, San Diego, CA, USA) or 1:1 ratio with AddaS03 (AddaS03 group) in 30 μl final injection volume (Invivogen, San Diego, CA, USA). Mice were injected intramuscularly at days 0, 14, and 42. Pre-vaccination serum was collected at day 0, test bleeds at day 28 and post vaccination sera (terminal bleeds) at day 56. Sera were collected after centrifugation of the samples at 5000 g for 15 minutes. Sera were heat-inactivated by incubation at 56°C for 30 minutes and stored in aliquots at −80°C until use.

### RBD ELISA

Microtiter plates were coated with RBD antigen (0.5 μg per well) in PBS and blocked in 5% bovine serum albumin (BSA) in PBS. Antisera from mice were diluted in PBST and added to the plates for 1 h (50 μl/well) in triplicate. RBD-specific antibodies from mouse antisera were detected by a horseradish peroxidase-conjugated goat anti-mouse secondary antibody (1:10,000; Cedarlane Laboratories, Burlington, ON, Canada) and peroxidase substrate (KPL, Gaithersburg, Md, USA). Absorbance was read at 450 nm. Absorbance values from two independent experiments are expressed as a percentage of the maximum OD450 signal ± SEM.

### RBD-ACE-2 binding assay

Microtiter plates were coated with RBD (1 μg/ml) in 0.1 M bicarbonate buffer overnight at 4 °C, washed three times with PBS + 0.05% Tween20, blocked with 2% BSA (in PBS + 0.05% Tween20) for 2 hour at room temperature and washed one additional time in PBS +0.05% Tween20. Pre- (pooled) or post- vaccination mouse serum was diluted in 3-fold (1:250-1:20250) with dilution buffer (PBS with 0.5% BSA, PBS, 0.05% Tween 20). The diluted sera (100 μl) were added for 30 minutes followed by addition of recombinant FLAG-tagged ACE-2 (100 μl at 400 ng/ml) for addition 2 hours. The final concentration of ACE-2 is 40 ng/well and the sera dilution is between 1:500-1:40500 after mixing (1:2 dilution). Plates were washed (3X) with PBS +0.05% Tween20 and bound FLAG-ACE-2 was detected with HRP-conjugated anti-FLAG antibody (1:20,000, Sigma cat# A8592) and peroxidase substrate (KPL, Gaithersburg, Md, USA). Absorbance was read at 450 nm and results plotted as % inhibition of control (no serum) and expressed as the reciprocal dilution that resulted in 50% inhibition (IC50).

### Psuedoparticle neutralization

HIV based pseudotyped virus with SARS-CoV-2 spike (CoV2pp), SARS-CoV-1 (CoV1pp) or glycoprotein of VSV (VSVpp) encoding a luciferase reporter were generated based on method described for HCVpp ^33^. The plasmid encoding the full length spike of SARS-CoV-2 with the terminal 19 amino acids deleted in order to increase yield ^31^ was generously provided by Dr. Bieniasz (The Rockefeller University). Variants containing mutations N501Y/E484K/K417N were constructed using standard molecular cloning techniques. The synthetic DNA fragment (Integrated DNA technologies Inc., Coralville, Iowa) containing corresponding mutations were used to replace the BamHI and AgeI fragment of pSARS-CoV-2Δ19 and confirmed by DNA sequencing. The plasmid encoding SARS-CoV-1 spike was commercially sourced (Sino Biologicals Cat# VG40150-G-N). For neutralization assays, 293T ACE-2 cells were plated on poly-lysine-coated 96-well plates one day prior to infection. CoV2pp was premixed with heat-inactivated diluted sera for 1 h at 37°C, followed by addition to 293T ACE-2 cells. The antibody-virus inoculum was replaced with fresh culture medium eight hours post-infection and cells processed 48 h post-infection with Nano-Glo luciferase assay system (Promega, Madison, WI). Luminescence was measured using an Enspire plate reader (PerkinElmer) and percentage virus entry (% Entry) calculated as follows: (Test sera luminescence signal/ PBS control luminescence signal) x 100. For IC50 titers, three fold dilutions of sera (1:50 to 1:109,350) were examined and IC50 titer expressed as the reciprocal of the serum dilution that resulted in a 50% reduction in virus entry.

### Live virus neutralization

SARS-CoV-2 (SARS-CoV-2/CANADA/VIDO 01/2020) was a kind gift from from Dr. Darryl Falzarano (Vaccine and Infectious Disease Organization). Both B.1.1.7 and B.1.351 were isolated from nasopharyngeal swabs by culture in Vero TMPRSS-2 cells. SARS-CoV-2 stocks were made and titers were determined in Vero E6 cells. Neutralizing antibody analysis was performed using a microneutralization assay based on the cytopathic effect (CPE) of SARS-CoV-2 on Vero E6 cells ^34^. Heat inactivated mouse sera samples were 2-fold serially diluted from dilution of 1/50 in infection medium. 100 plaque forming unit (PFU) of SARS-CoV-2 was then added, and 96 well plates were incubated for 1 hour at 37°C. At the end of the incubation, the mixture was transferred onto 96-well microtiter plates pre-seeded overnight with cells. Plates were incubated for 3 days at 37°C and terminated by fixing in formaldehyde, followed by staining with crystal violet. Cytopathic effect (CPE) was then quantified and the N100 microneutralization titer was defined as the reciprocal of the highest sample dilution that protects from CPE. If no neutralization was observed, samples were arbitrarily assigned a N100 titer value of 25 (half the minimum dilution).

### T cell assay

Immediately after euthanasia, mouse spleens were extracted and transferred to culture media. Splenocytes were isolated and red blood cells lysed with RBC Lysis Buffer (BioLegend, CA, USA). Splenocytes from each vaccination group were pooled (4 or 3 spleens per pool, 2 pools per group) and dispensed in triplicate to 96-well round bottom plates (Corning, NY, USA) for analysis of test groups: 1) negative control consisting of media alone; 2) Non-specific peptides consisting of a pool of 55 peptides spanning hepatitis-C genotype 1a H77 E1E2 glycoproteins; and 3) Specific RBD peptides consisting of a pool of 66 peptides (15 amino acids each) spanning the SARS-CoV-2 Spike RBD 319-591 region with 11 amino acid overlap. Both the negative and peptide groups had DMSO added to match the concentration in the RBD peptide pool group (0.4% v/v). After a 1.5 hr incubation at 37°C with 5% CO2, Brefeldin A and Monensin (Biolegend, CA, USA) were added, followed by an additional 5 hr of incubation. Cells were then centrifuged, washed with PBS, and stained for dead/live (Biolegend), surface markers (CD3, CD4, and CD8), intracellular cytokines (IFN-γ and TNF-α). FACS Analysis was performed using Fortessa-SORP flow cytometer analyzer (BD Biosciences, CA, USA) flow cytometer and analyzed with FlowJo.

## Results

### Purification of recombinant SARS-CoV-2 RBD

Most neutralizing antibodies target the SARS-CoV-2 RBD ^20^ and this can be expressed at high levels in transfected mammalian cells. Therefore, we developed the RBD of the spike protein (amino acid residue 319-591) as our vaccine antigen ^9^. To streamline future clinical development, we used our previous strategy for expression of the HCV glycoprotein vaccine ^33^. An N-terminal TPA signal peptide sequesters the RBD for secretion and downstream of the RBD, a C-terminal HRV 3C recognition site followed by a monomeric FC (mFc) tag is used to facilitate purification (Figure S1). For pre-clinical studies, a lentivirus based vector constitutively expressing RBD-mFc in CHO cells was used ^32^. The RBD-mFc was purified from cell culture media with a protein G based column (Figure S1b) and was used to immunize mice either alone or adjuvanted with either Alum containing the TLR4 agonist MPL or a tocopherol-containing emulsified adjuvant (AddaSO3) shown previously to be of value in pandemic vaccines ^35,36^. Mice (CB6F1) were immunized with 1 μg and then boosted twice on days 14 and 42.

### RBD vaccine induced seroconversion and T cell responses

We examined antibody titers 2 weeks after the second immunisation (Day 28, test bleed) and after the third immunisation (Day 56, final bleed) (Figure 1). To determine the immunogenicity of our vaccine, we examined vaccinated mouse sera for RBD binding (Figure 1). As expected, RBD formulated with or without adjuvant successfully induced seroconversion in mice, however the titer of RBD binding antibodies was much higher when RBD was formulated with either Alum/MPLA or AddaS03 adjuvants (Figure 1). Using the final bleed sera for analysis, we detect significantly higher RBD binding antibody titers (up to 1/80,000 dilution) in either adjuvanted RBD as compared with RBD alone. Since we found that antibody titers were highest after the third immunisation, we focused our further analyses on samples from the final bleed.

**Figure 1.**
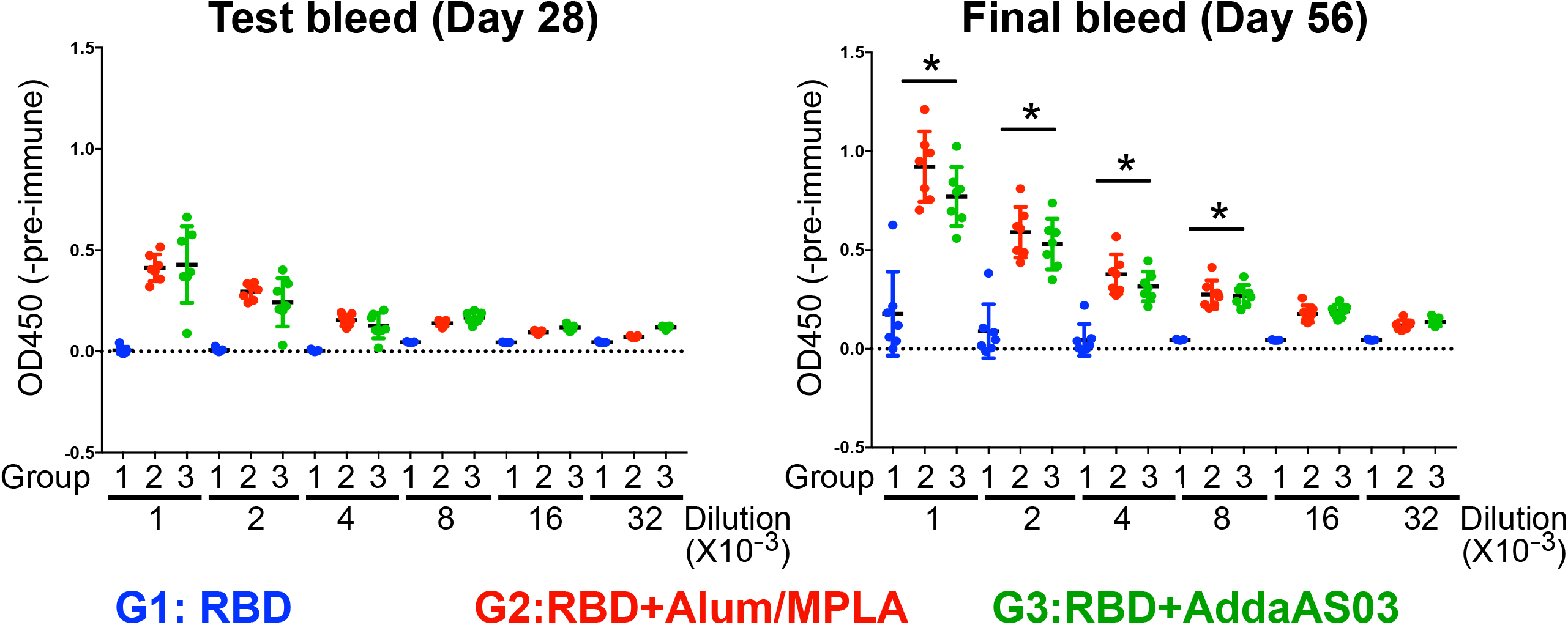
RBD-specific antibody titers following vaccination in test-bleed (D28) and in final bleed (D56). Recombinant RBD (319-591) of SARS-CoV-2 coated plates were probed with pre-or post-vaccination mouse antiserum (test and final bleed) and bound RBD-specific antibodies detected by horseradish peroxidase-conjugated anti-mouse secondary antibody and peroxidase substrate. RBD binding activity of post-vaccinated antiserum between 1/10,000 to 1/320,000 dilution are shown. The optical densities at 450 nm subtracted from pre-immune control (OD450-Pre-immune sera) (mean ± SEM) are measured. (*) indicates p<0.05 in Tukey’s multiple test between G1 and G2/G3. G1, RBD (blue); G2, RBD/Alum+MPLA (Red); G3, RBD/AddaS03 (Green). Lighter color symbols represent the test bleed sera and the darker color symbols represent the final bleed sera.

To assess T-cell responses in vaccinated mice, we examined the production of TNF-α and IFN-γ from RBD-specific T cells. We stimulated splenic cells from immunized mice with either overlapping SARS-CoV-2 RBD peptides or HCV E1/E2 peptides as a control, and then examined the production of either TNF-α or IFN-γ in CD4+ and CD8+ cells. We found that vaccination with RBD was capable of inducing robust CD4+ and CD8+ T cell responses (Figure 2 and S2). A higher proportion of CD4+ or CD8+ T-cells produced TNF-α, IFN-γ, or a combination of both in mice that were vaccinated using RBD adjuvanted with either Alum/MPLA or AddaS03. Both CD4+ and CD8+ cells were activated in response to RBD peptides (black bars), but not in response to HCV E1/E2 peptides (grey bars), demonstrating that the T-cell responses were specific for SARS-CoV-2 RBD. The high levels of production of TNF-α and IFN-γ by RBD-specific CD4+ and CD8+ T cells indicate a strong TH-1 response. The TH-1 response is important for developing cytotoxic T cell (CTL) immunity. Altogether, our data show that vaccination with RBD formulated with Alum/MPLA or AddaS03 elicits high levels of RBD binding antibodies along with a robust cellular immune response.

**Figure 2.**
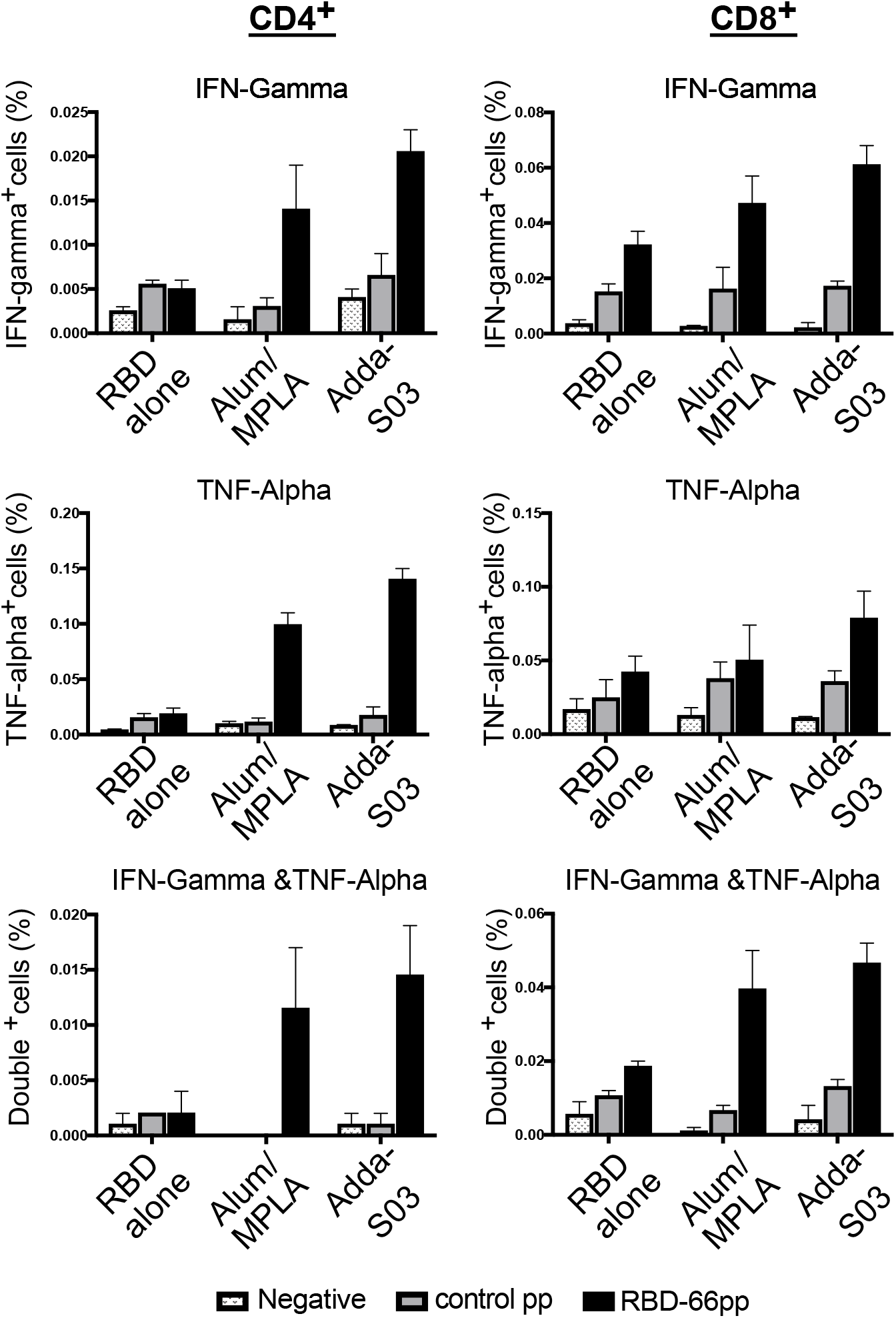
Activation of RBD-specific CD4+ and CD8+ T-cells following vaccination. Splenocytes from vaccinated mice were stimulated *in vitro* and intracellular production of cytokine (and/or IFN-γ and/or TNF-α) was detected by multi-color flow cytometry. The percentage of CD4+ and CD8+ T cells that are expressing IFN-γ, TNF-α or both are shown, left panels showing CD4+ T cells and right panels showing CD8+ T cells. Control-55pp represents splenocytes that are stimulated with a pool of 55 peptides spanning HCV E1E2; RBD-66pp represents splenocytes that are stimulated with a pool of 66 peptides spanning SARS-CoV-2 Spike RBD 319-591 (see methods and materials). Splenocytes from each vaccination group were pooled into two groups and the average of these two groups are shown. Dot plots of a representative experiment are shown in figure S2.

### Antisera blocks RBD binding to ACE-2

One possible mechanism of neutralization by RBD specific antibodies is to interfere with the interaction between the host receptor, ACE-2 and the spike protein. We developed an ELISA based RBD-ACE-2 binding assay and examined the effect of pre- or post-vaccination mouse sera on the interaction between ACE-2 and RBD. We observed dose-dependent inhibition of RBD binding to ACE-2, by antisera from mice immunized with RBD formulated with either Alum/MPLA or AddaS03 (Figure 3a). Pooled pre-immune sera did not inhibit RBD-ACE-2 interaction. An IC50 reciprocal titer for each mouse serum was calculated and values of adjuvanted RBD ranged between 1298 to 4150. When we compared the effect of adjuvants, the IC50 values from the antisera of mice immunized using Alum/MPLA formulation were higher than those immunized using AddaS03 formulation (Figure 3b).

**Figure 3.**
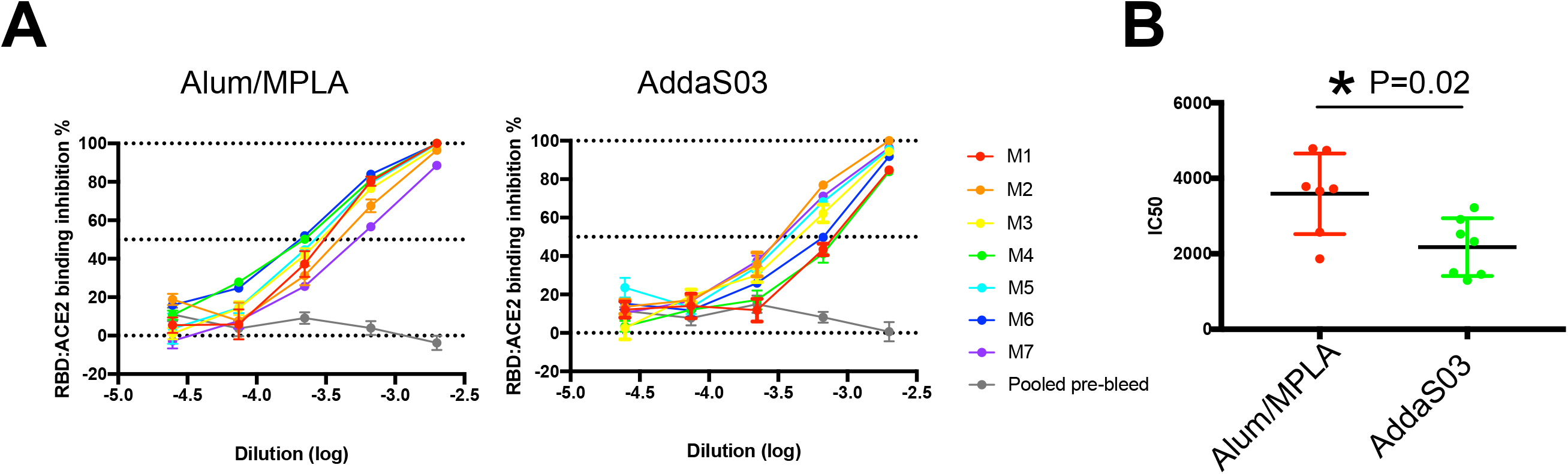
Vaccinated antisera blocks RBD-ACE-2 interaction. (A) 3-fold serial diluted antisera was added to micro-titer plates coated with recombinant RBD protein. After 30 minutes incubation, FLAG-tagged ACE-2 protein were added and detected with anti-FLAG antibody. Pooled pre-immune serum of each group is used as control. Amount of bound ACE were determined. Each colored line represent serum of a mouse and pooled pre-immune serum is in grey. (B) Reciprocal Inhibitory Dose 50 (IC50) was calculated for sera of each animals. Comparison of IC50 between the two adjuvants is show. (*) indicates p<0.05 in Mann Whitney test.

### Antisera from vaccinated mice neutralise infection*in vitro* by parental and variant SARS-CoV-2 strains

We also examined inhibition of SARS-CoV-2 infection *in vitro* by antisera using both lentivirus based SARS-CoV-2 pseudoparticles (PP) (Figure 4a) and infectious SARS-CoV-2 virus (Figure 4b). Consistent with our previous observations, antiserum from mice vaccinated using RBD formulated with Alum/MPLA or AddaS03 neutralized SARS-CoV-2pp infection of 293 cells expressing ACE-2 whereas pre-immune serum did not. SARS-CoV-2 pp entry was prevented by antiserum from all 14 mice when diluted 1:100, whereas 6/7 similarly diluted sera from mice that were vaccinated with unadjuvanted RBD alone did not completely prevent SARS-CoV-2pp entry (Figure 4a). Furthermore, we examined the neutralization of infection of Vero E6 cells by the intact SARS-CoV-2 virus. Infection of Vero E6 cells by SARS-CoV-2 results in cytopathic effects (CPE). We used a microneutralization assay in which 100 plaque forming units (PFUs) was incubated with serially diluted sera prior to infection of Vero E6 cells ^37^. The greatest dilution that prevents cell lysis is the N100 neutralization titer (See Materials and Methods) ^34^. Antisera from mice vaccinated with RBD formulated with either Alum/MPLA or AddaS03 had greater virus-neutralizing activity than pre-immune controls (Figure 4b). The titer of these vaccinated sera were 18 fold higher than sera obtained from patients convalescent from COVID19 infection ^37^, indicating that adjuvanted RBD elicits an excellent humoral response.

**Figure 4.**
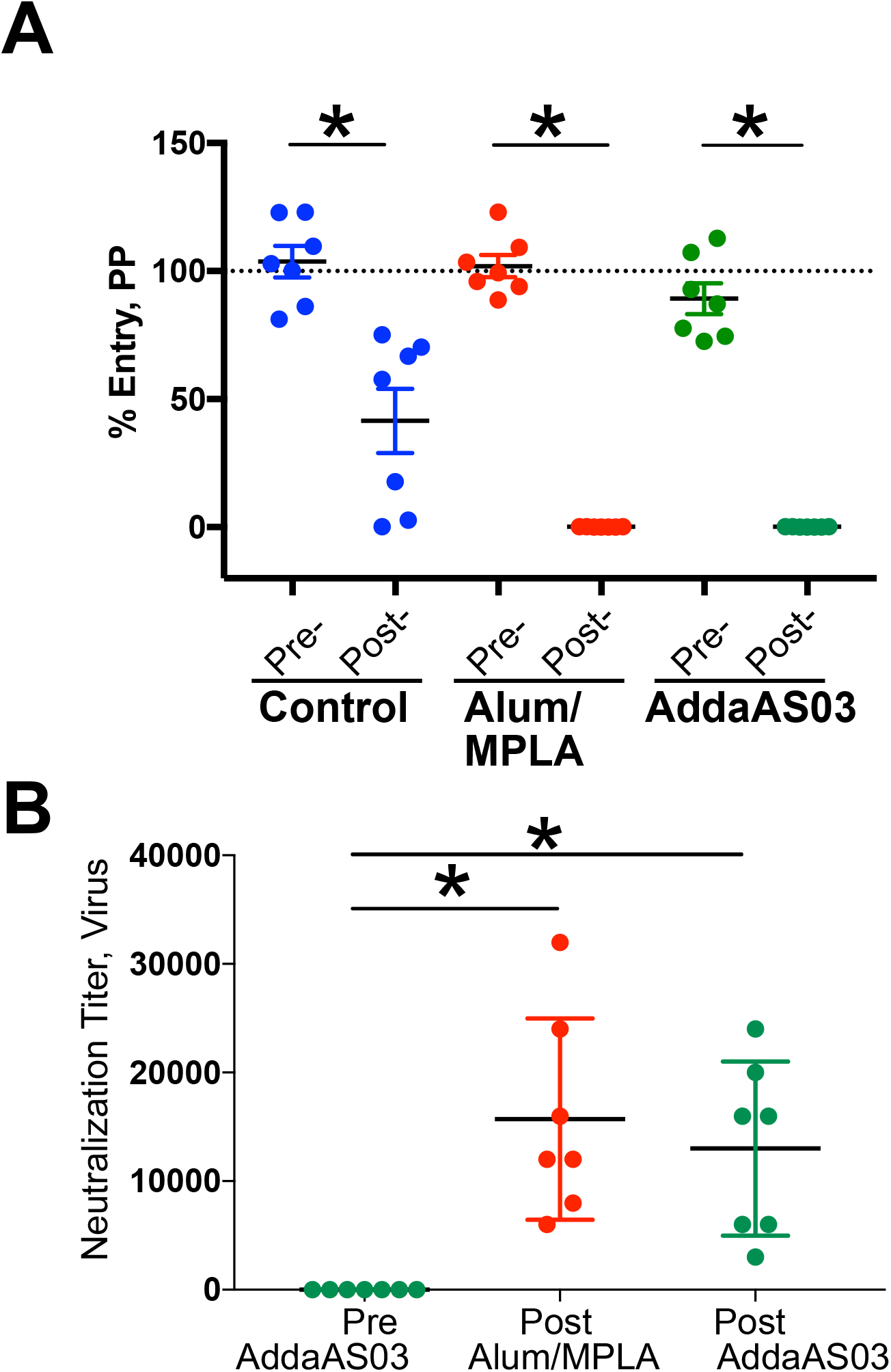
Vaccination-induced neutralizing antibodies (nAb) protects from SARS-CoV-2 infection. Pre-immune or post-vaccinated mice sera were evaluated for their ability to neutralize SARS-CoV-2 pseudoparticles (CoV2pp) (A) or infectious SARs-CoV-2 virions (B). (A) Neutralization CoV2pp was performed using pre- and post-vaccination sera (1:50) in 293T ACE-2 cells. The group means in triplicates with SEM were plotted and the level of entry was normalized to entry of CoV2pp in the presence of PBS as 100%. (*) indicates p<0.05 by two-way ANOVA and Tukey’s multiple comparison test. (B) Neutralization titer (N100) of vaccinated mouse sera was determined in Vero E6 cells using infectious SARS-CoV-2. Serially diluted sera were pre-incubated with 100 PFU of SARS-CoV-2 for 1 hour at 37°C followed by addition to Vero E6 cells. Three days post-infection, cells were formaldehyde fixed and stained with crystal violet. Neutralization titer was determined as the minimal dilution of each mouse serum required to prevent CPE.

Variants of SARS-CoV-2 have arisen in the general population during the past 6 months, and two of particular concern are B1.1.7 and B.1.351 originating in the UK and South Africa respectively. These variants are spreading worldwide. Variant mutations within S affect its interaction with the host receptor ACE-2 ^38^. Variant B.1.351 in particular has been shown to reduce vaccine neutralization *in vitro* ^39^. Here, we focus on three mutations (K417N, E484K and N501Y) found within RBD of the B.1.351 (also found in P.1 variants, originating in Brazil) ^40^. We engineered these three mutations into SARS-CoV-2pp, then examined the ability of serially diluted post-vaccination sera or pooled pre-immune sera to neutralize variant or wild type (WT) pp entry (Figure 5a). We observed a dose dependent effect on neutralization (Figure S3) and determined the IC50 value for antisera from each mouse against either the WT or variant SARS-CoV-2pp. Variant neutralization IC50 values were not significantly different from WT IC50 neutralization (Figure 5a).

**Figure 5.**
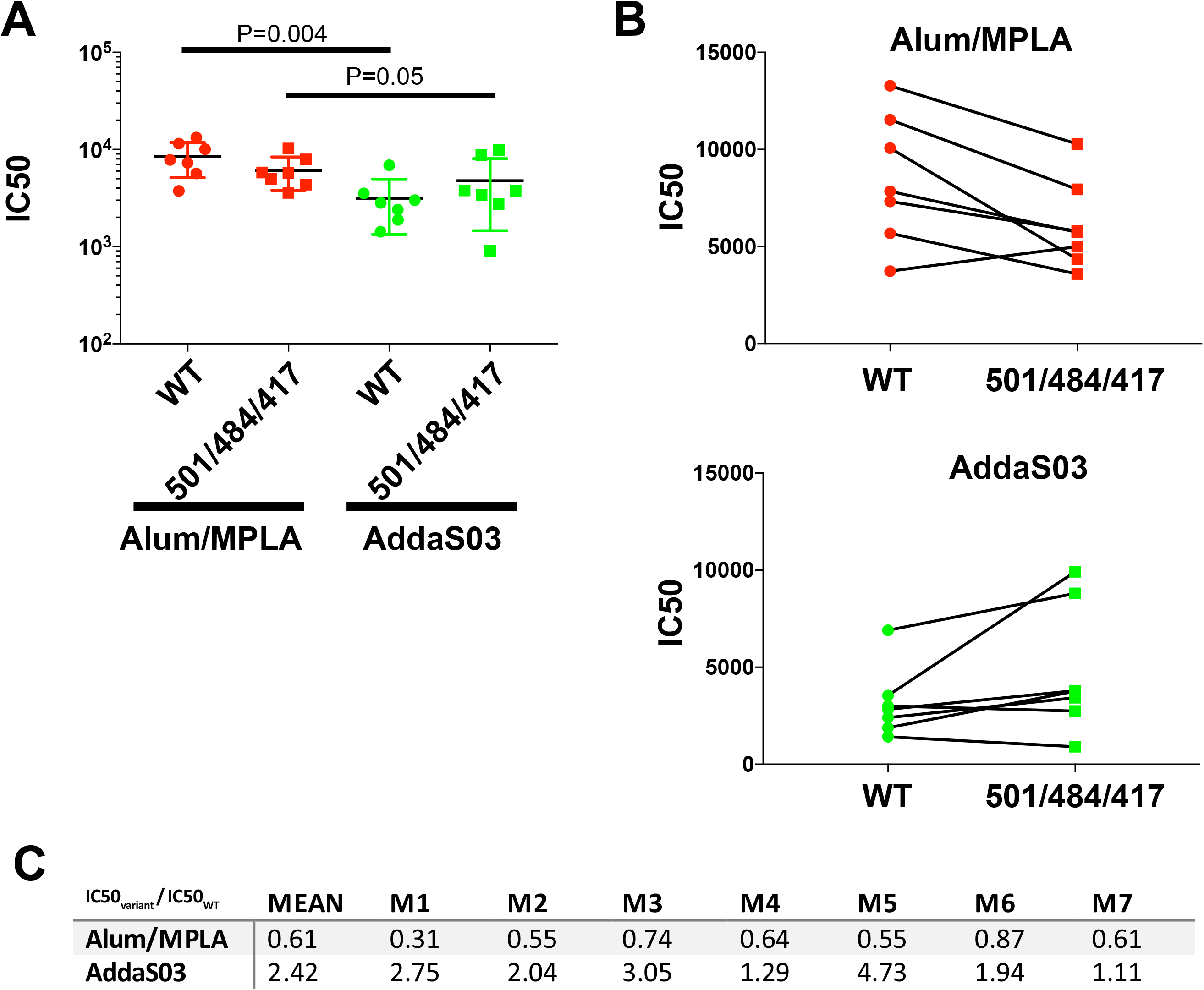
Antisera exhibits similar neutralization activity against SARS-CoV-2pp containing variant mutations N501Y/E484K/K417N. Neutralization activity of mouse sera were tested at 1:50 to 1:109,350 in 3 fold dilution against lentivirus particles pseudotyped with either spike of SARS-CoV-2(WT) or S encoding triple mutations N501Y/E484K/K417N. Pre- or Post-vaccination sera were pre-incubated with pseudoparticles (PP) encoding different surface proteins for 1 hour followed by addition to 293T ACE-2 cells. 48 hour post-transduction, entry of PP were quantitated by luciferase activity. Result are done in duplicate normalized to entry of PP in the presence of PBS. IC50 of neutralization activity was calculated and compared between groups. (A) mean and standard error of mean of IC50 is shown. significant P-values by two-way ANOVA and Tukey’s multiple comparison test were shown. (B) Change of neutralization (IC50) between WT and variant (501/484/417) PP were shown. Representative of two independent experiments are shown. (C) Comparison of IC50 between WT and variant are shown. The ratio of IC50variant /IC50WT for each individual mouse and mean for each group are calculated. This is result showing average of two independent experiments done.

Consistent with what we saw in the RBD-ACE2 binding assay, there was a small reduction in the neutralization IC50 value by antisera of mice vaccinated with RBD adjuvanted with AddaS03 compared to those vaccinated with Alum/MPLA (Figure 5a). In the group vaccinated with RBD formulated with Alum/MPLA the neutralization IC50 of the variant was 1.6 fold lower than WT whereas in the group vaccinated with AddaS03 the IC50 was 2.4 fold higher than WT (Figure 5b and 5c). Nonetheless, the efficacy of neutralizing both WT and variant SARS-CoV-2 is comparable within either adjuvant (Figure 5b). Interestingly, the fold change in neutralization of WT and variant SARS-CoV2pp appears to somewhat depend on the adjuvant.

Besides the three mutations (K417N, E484K and N501Y), variant B 1.1.7 or B 1.351 each has additional mutations outside of RBD (10 mutations in B 1.1.7 and 12 mutations In B 1.351) ^40^. In order to investigate whether these additional mutations affect the sensitivity of these variants to neutralization, we tested the neutralization of these variants using infectious virus isolated from clinical samples. In Figure 6, the antisera from mice vaccinated with either RBD Alum/MPA or RBD AddaS03 neutralized the variant B 1.1.7 with the same efficiency as WT virus. However, we observed a statistically significant reduction in the ability of sera to neutralize the B 1.351 variant vs WT. The mean N_100_ titer for neutralizing the B.1.351 variant was 1.6 fold or 2.5 fold lower than that of the WT virus when RBD formulated with Alum/MPLA or AddaS03, respectively (Figure 6).

**Figure 6.**
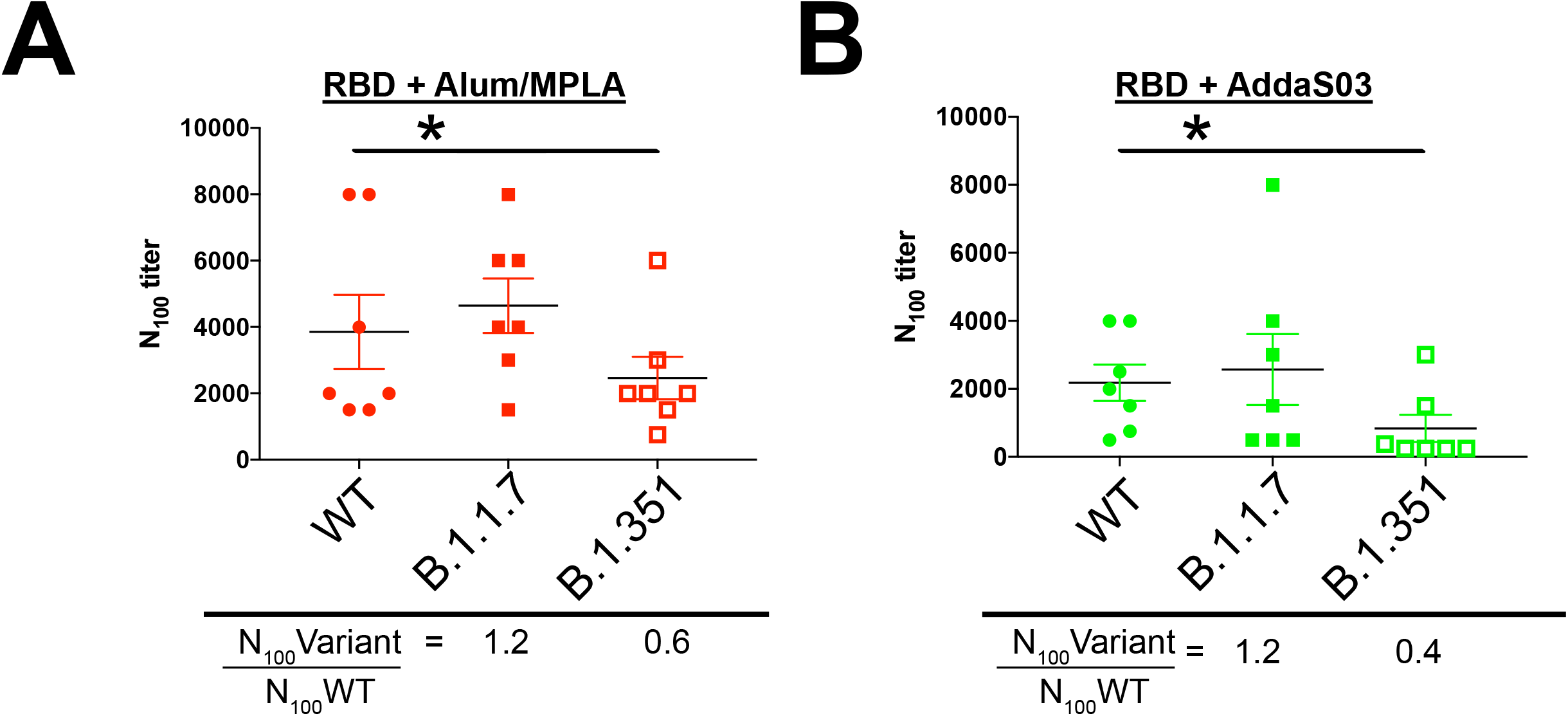
Antisera exhibits neutralization against SARS-CoV-2 B.1.1.7 and B.1.351 variants. Neutralization titer (N100) of vaccinated mouse sera was determined in Vero E6 TMPRESS-2 cells using infectious WT SARS-CoV-2, B 1.1.7 or B 1.351 variant. Serially diluted sera were pre-incubated with 100 PFU of SARS-CoV-2 for 1 hour at 37°C followed by addition to Vero E6 TMPRESS-2 cells. Three days post-infection, cells were formaldehyde fixed and stained with crystal violet. Neutralization titer was determined as the minimal dilution of each mouse serum required to prevent CPE. Comparison of IC50 between WT and Variant are shown. The mean ratio of IC50variant /IC50WT for each group are calculated. This is result showing average of two independent experiments done. * indicates significant P-values by two-way ANOVA and Tukey’s multiple comparison test between WT and B 1.351.

We further examined the ability of this vaccine to block SARS-CoV-1 infection (Figure 7b). Compared to SARS-CoV-2 (Figure 7a), neutralisation of SARS-CoV-1 pp was apparent but significantly reduced at a 1:500 dilution. Protection was specific because the antisera did not confer any protection to VSV pp (Figure 7c).

**Figure 7.**
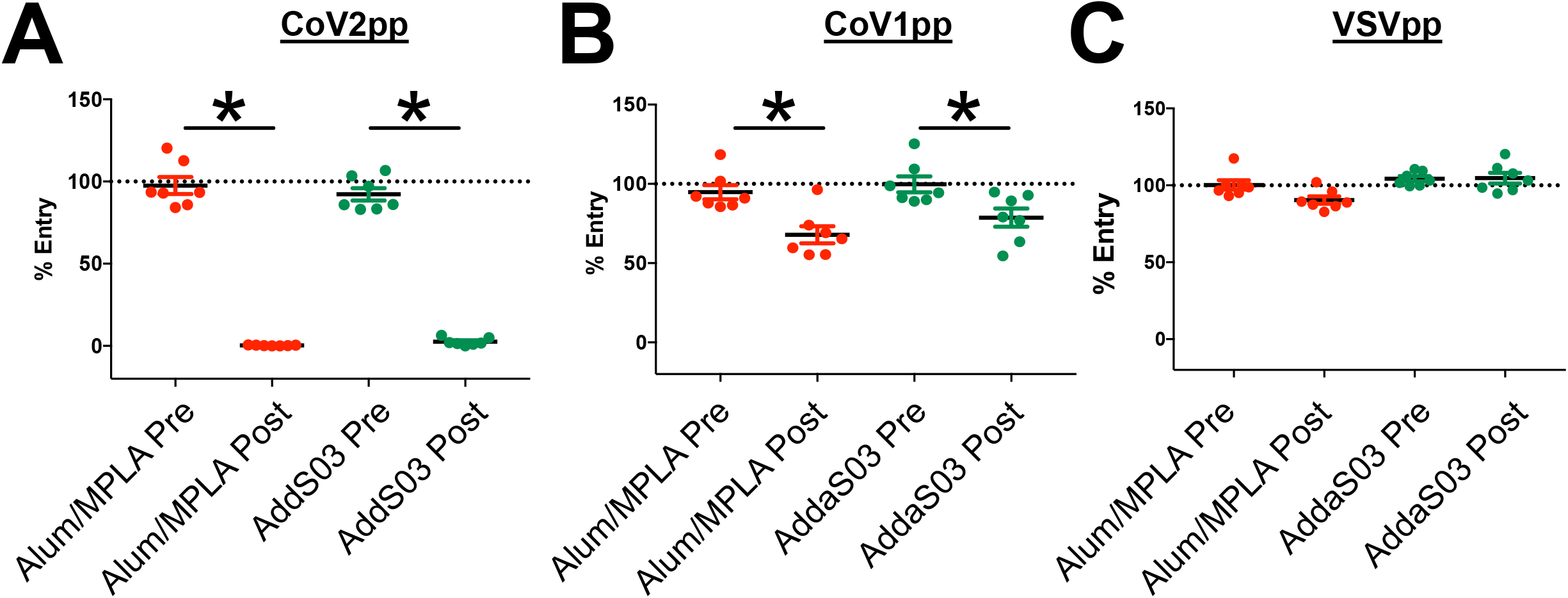
Antisera from RBD SARS-CoV-2 vaccinated mice exhibits cross-neutralization against SARS-CoV-1. Neutralization activity of mouse sera were tested at 1:500 against pseudo-typed virus particles (PP) expressing either SARS-CoV-2 spike (CoV2pp) (A), SARS-CoV-1 spike (Cov1pp) (B) or the glycoprotein of VSV (VSVpp) (C). Pre- or post-vaccination sera were pre-incubated with PP and added to 293T ACE-2 cells and assessed for luciferase activity according to the Materials and Methods. Triplicate samples were normalized to entry of PP control (PBS) and the mean with SEM plotted. (*) indicates p<0.05 by two-way ANOVA and Tukey’s multiple comparison test. A representative of three independent experiments is shown.

## Discussion

A number of COVID19 vaccines based on the spike protein are currently being deployed throughout the world, however there are rising concerns about the ability of vaccines to protect from infection by the variant strains of SARS-CoV-2. We have evaluated adjuvanted recombinant spike RBD of SARS-CoV-2 for use as a prophylactic vaccine. In this study, the formulated RBD with either Alum/MPLA or AddaS03 adjuvants was immunogenic in mice and induced robust, specific CD4+ & CD8+ T responses. Many studies have reported reduction of neutralization against B 1.351 ^4–6^. Most of them have characterized antisera from clinical trials using mRNA based vaccines. The observed reduction in IC50 was 4-12x ^6,41^, whereas the reduction using our RBD vaccine was somewhat less than 3 fold. However, since the mRNA antisera was derived from humans and our adjuvanted vaccine was derived from mice, it remains to be seen if our results can be extrapolated to humans. If they can, then our vaccine could be most effective against the known SARS-CoV-2 variants. In addition, adjuvanted subunit proteins have been proven to be very effective and very safe in protecting against many viral infections including the hepatitis B virus, hepatitis A virus, human papilloma virus, varicella zoster virus, and many other viruses. Therefore, our approach offers potential advantages in dealing with SARS-CoV-2 variants effectively and with a greater safety profile as compared with newer technologies that have been shown to display specific toxicities, albeit rarely ^42–45^. In addition, ready scale-up of our vaccine process to meet global demands for booster shots is feasible.

Currently, the duration of protection conferred by vaccination has not been fully determined. However, the half-life of anti-SARS-CoV-2 antibodies in convalescent serum has been reported to be approximately 49 days ^19^. It is also unclear whether declining antibodies equates with a lack of protection. It has been reported that long lived memory plasma cells are generated by SARS-CoV-2 infection and strong antibody responses to SARS-CoV-1 have been reported from some individuals 17 years after infection ^46^. Adjuvants including TLR7/8 agonists have been shown to generate more durable humoral responses in non-human primates against HIV ^47^. Recent work in non-human primates using nanoparticles containing RBD formulated with various adjuvants has indicated that extension of protection is possible ^25^. Further characterization of the immune response in both convalescent SARS-CoV-2 patients and vaccinees is needed to determine whether boosters will be required to provide long lasting protection. Considering the worldwide effort to immunize against SARS-CoV2, a prolonged response after the primary vaccination is highly desirable in order to reduce the requirement for subsequent booster immunizations.

From the work of others, recombinant SARS-CoV-2 spike RBD antigen has produced mixed results when utilized as a vaccine antigen. In one study, RBD (expressed and purified from Sf9 insect cells) conferred protection in rhesus macaques ^23^. On the other hand, a separate report showed that RBD (expressed and purified from mammalian cells) was poorly immunogenic in mice ^48^. However, when full-length S protein was used to prime followed by a boost with RBD, neutralizing antibodies were generated ^48^. Different cell types have been used to produce either recombinant S or RBD antigen, such as yeast, plant and insect cells ^23,49,50^. However production in host mammalian cells (in our study, CHOK1) may result in producing a RBD antigenic domain closely resembling that generated during virus infection in human cells (including post-translational modifications such as glycosylation and correct folding) ^38^. Along with promising protection data reported in rhesus macaques ^23^, our studies further encourage clinical development of an adjuvanted recombinant RBD vaccine. In addition to the vaccine antigen, formulation of the antigen may require further optimization. During development of an mRNA based vaccine of Pfizer, the choice between full length S (BNT162b2) and RBD (BNT162b1) was determined by the breadth of T cell response beyond epitopes within RBD, thus prompting the decision to use full length spike as the vaccine antigen ^26–28^.However, the relative role of antibodies or T cell responses in protection from COVID19 remain unknown although based on all other viral vaccines, neutralizing antibodies likely play a central role in protection. In our hands, yields of recombinant RBD were much higher than that of full-length S, an important factor in delivering the vaccine to global populations. In this regard, we have developed a GMP-grade CHO cell-line directly expressing the RBD domain itself which is available from us for clinical development around the globe.

Given the concerns about the ability of the current vaccines to protect from infection by variant strains of SARS-CoV-2, we were particularly encouraged by the strong cross-neutralisation of SARS-CoV-2pp encoding dominant mutations (501/484/417) found in B.1.351, and P.1 variants. Furthermore, we observed evidence of cross-neutralisation of B1.1.7 and B.1.351 infectious variants, albeit with somewhat reduced neutralization activity against B.1.351. This is consistent with the functional constraints placed on the RBD domain such that they must maintain their interaction with the host receptor, ACE-2 for cell entry of the virus. Even for viruses that exhibit high mutation rates and sequence variability (for example, HCV), viral epitopes that interact with host receptors are usually better conserved and less likely to tolerate mutations that inhibit interactions between the virus and its receptor ^51^. Currently, approved COVID19 vaccines employ full length S protein as the vaccine antigen. Although the RBD has been shown in several studies to be an immunodominant region of S ^19,20^, neutralizing epitopes in regions outside the RBD, such as the N-terminal domain (NTD) and C-terminal domain of S2 have been reported ^21,52^. These regions of S1 are more diverse amongst coronavirus strains and appear to be more readily mutated ^52^. For example, the B.1.1.7 variant was shown to be refractory to neutralization by NTD-specific monoclonal antibodies ^52^. However, there are also examples of escape mutations within the RBD itself ^41^. The cross neutralisation of variant strains that we observed in our study is consistent with induction of a broad polyclonal response to a multiplicity of RBD neutralizing epitopes. This is similar to the neutralization of SARS-CoV-2 variants by convalescent sera, whereas various isolated RBD-specific monoclonal antibodies may fail to provide neutralisation of virus infectivity ^53^.

Mutation of E484 in the RBD has shown to reduce antibody neutralization *in vitro* by sera from specific patients^54^. This suggests that a patient’s genetic background may affect their immune response and, perhaps, contribute to immune escape. Furthermore, the evolving strains of SARS-CoV-2 and their dominance in the population may require that vaccines match the circulating strain, similar to influenza vaccine strategies. For example, Cele et al. reported that variants of the South African lineage (B1.351) evolved to escape neutralization from convalescent sera that was collected earlier in the pandemic from the same region ^55^. Our data showing strong *in vitro* cross-neutralisation of different strains by our RBD antisera suggests that along with robust cellular immune responses, RBD-based vaccines could be of value in dealing with emerging variants.

Another approach to SARS-CoV-2 vaccination and its evolving variants is to focus on conserved SARS-CoV-2 epitope(s) in order to circumvent viral escape mutations. A similar approach to produce a universal influenza vaccine and/or broad HIV vaccine could be examined ^56,57^. For example, vaccination strategies that target the less immunogenic but more conserved S2 region of the SARS-CoV-2 S protein may be worthwhile. A similar strategy has shown promise with the hemagglutinin (HA) antigen of influenza virus, where broadly protective antibody responses targeting the conserved stalk region of HA were induced during a phase I clinical trial ^56^. Another approach, employed for protection against SIV, has been sequential immunizations of closely related SIV antigens which broadened the humoral response to SIV ^58^. Interestingly, there is one report showing that broad protection against many coronaviruses is possible. Co-presentation of multiple RBDs from several different coronaviruses on nanoparticles induced a broad cross-neutralisation and strong immune response in mice ^24^. Continued research on an adjuvanted RBD vaccine could be of value in our global response to this on-going pandemic and to prepare for future zoonotic infections by SARS-related coronaviruses. Our data showing cross-neutralization against SARS-CoV-1 pp variants indicates that there are significant albeit limited conservation of epitopes between the RBDs of these viruses. This is consistent with other studies showing cross-neutralising antibodies between SARS-CoV-1 and SARS-CoV-2^10^. If we are able to identify and optimize the response to the conserved epitopes, a universal vaccine against very diverse coronaviruses may be possible. Surveys of coronaviruses within bat caves in China reveal diverse reservoirs of many closely related yet distinct coronaviruses, some of which also use ACE-2 for virus entry ^59,60^. Continued research on developing a universal coronavirus vaccine could potentially prevent yet another zoonotic infection by coronaviruses.

## Acknowledgements

We thank Darryl Falzarano (Vaccine and Infectious Disease Organization), Frauke Muecksch and Paul Bieniasz (The Rockefeller University) for kindly providing valuable reagents; Darci Loewen-Dobler for technical assistance; Staffs of HSLAS at University of Alberta for animal work. Flow cytometry was performed at the University of Alberta Faculty of Medicine and Dentistry Flow Cytometry Facility, which receives financial support from the Faculty of Medicine and Dentistry and Canadian Foundation for Innovation (CFI) awards to contributing investigators. This work was supported by Canadian 2019 Novel Coronavirus (COVID-19) Rapid Research Grant (M.H. and D.L.T), Alberta Innovates Health Solutions, and the Western Economic Development Program of Alberta.

---

**Figure S1.**
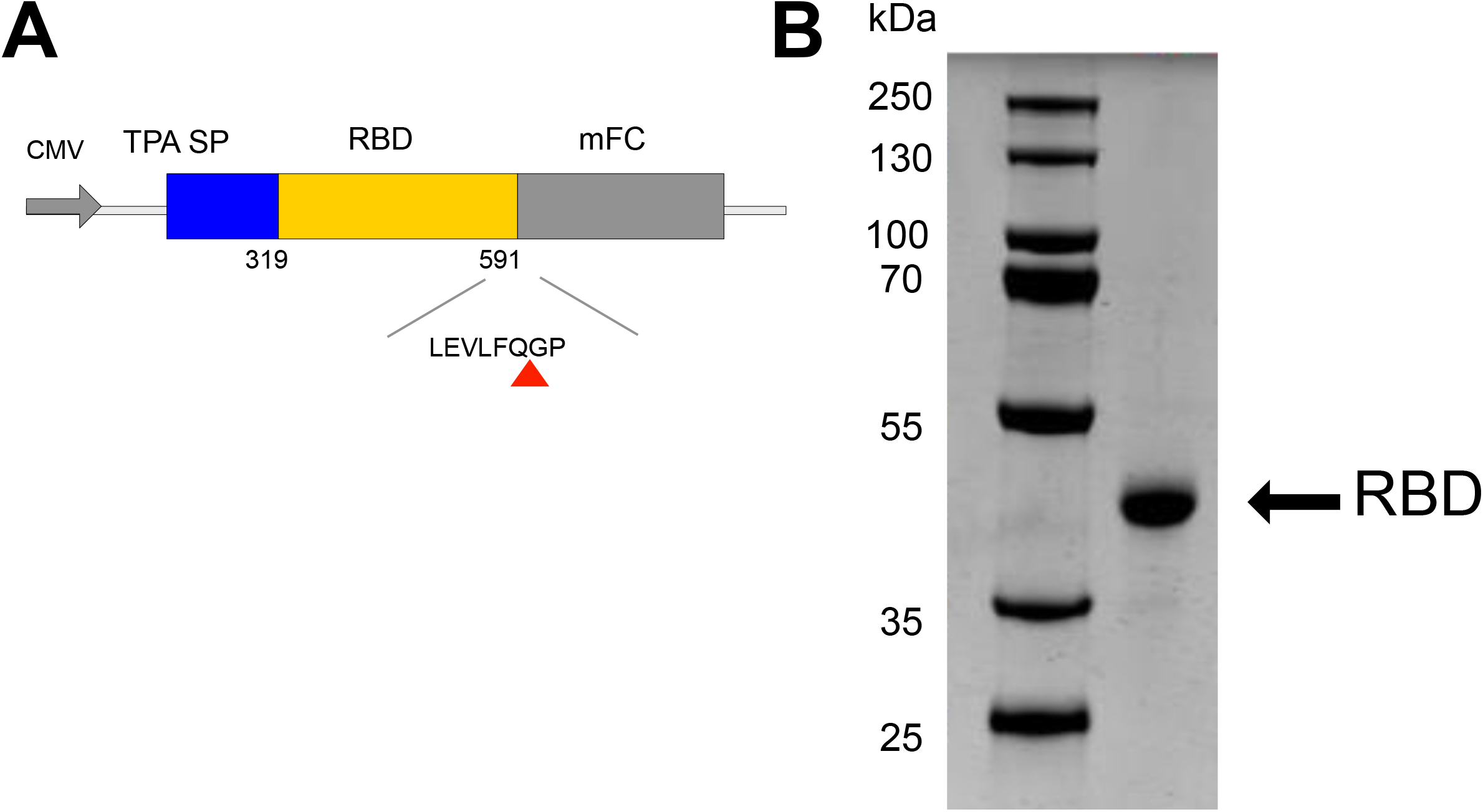
Purification of RBD from an mFc-tagged precursor. (A) Schematic representation of mFc-tagged RBD of SARS-CoV-2. (A) Amino acids 319-591 of the receptor binding domain of SARS-CoV-2(RBD (319-591) CoV2) (Genbank AAP13567.1) was expressed under the control of the CMV promoter (CMV) and preceded by the signal sequence from tissue plasminogen activator (TPA). A C-terminal human monomeric IgG1 Fc tag (mFc) was inserted downstream of the RBD that contained a human rhinovirus 3C (HRV3C) protease cleavable linker (LEVLFQGP). Red triangle indicates cleavage site by HRV3C protease. (B) RBD (319-591) CoV2 (2μg load) was purified from the mFc tagged precursor according to the Methods & Materials, separated by reducing SDS-PAGE and stained with Coomassie brilliant blue G250.

**Figure S2.**
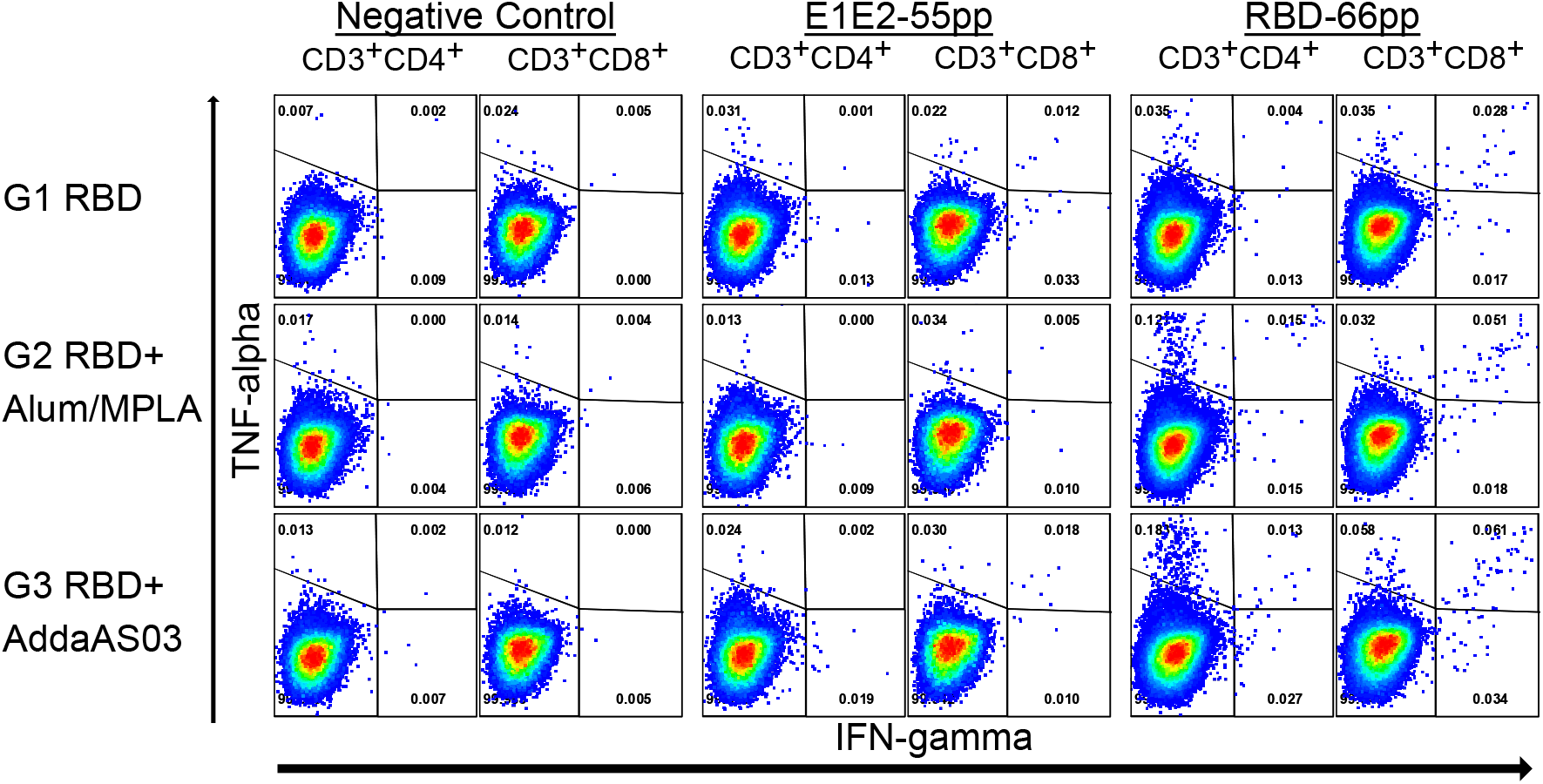
Activation of RBD-specific CD4+ and CD8+ T-cells following vaccination. Splenocytes from vaccinated mice were stimulated *in vitro* and intracellular production of cytokine (IFN-γ and/or TNF-α) was detected by multi-color flow cytometry. Dot plots of a representative experiment are shown. Numbers indicate the percentage of CD4+ and CD8+ T cells that are expressing IFN-γ, TNF-α or both. E1E2-55pp represents splenocytes that are stimulated with a pool of 55 control peptides spanning HCV E1E2; RBD-66pp represents splenocytes that are stimulated with a pool of 66 peptides spanning SARS-CoV-2Spike RBD 319-591.

**Figure S3.**
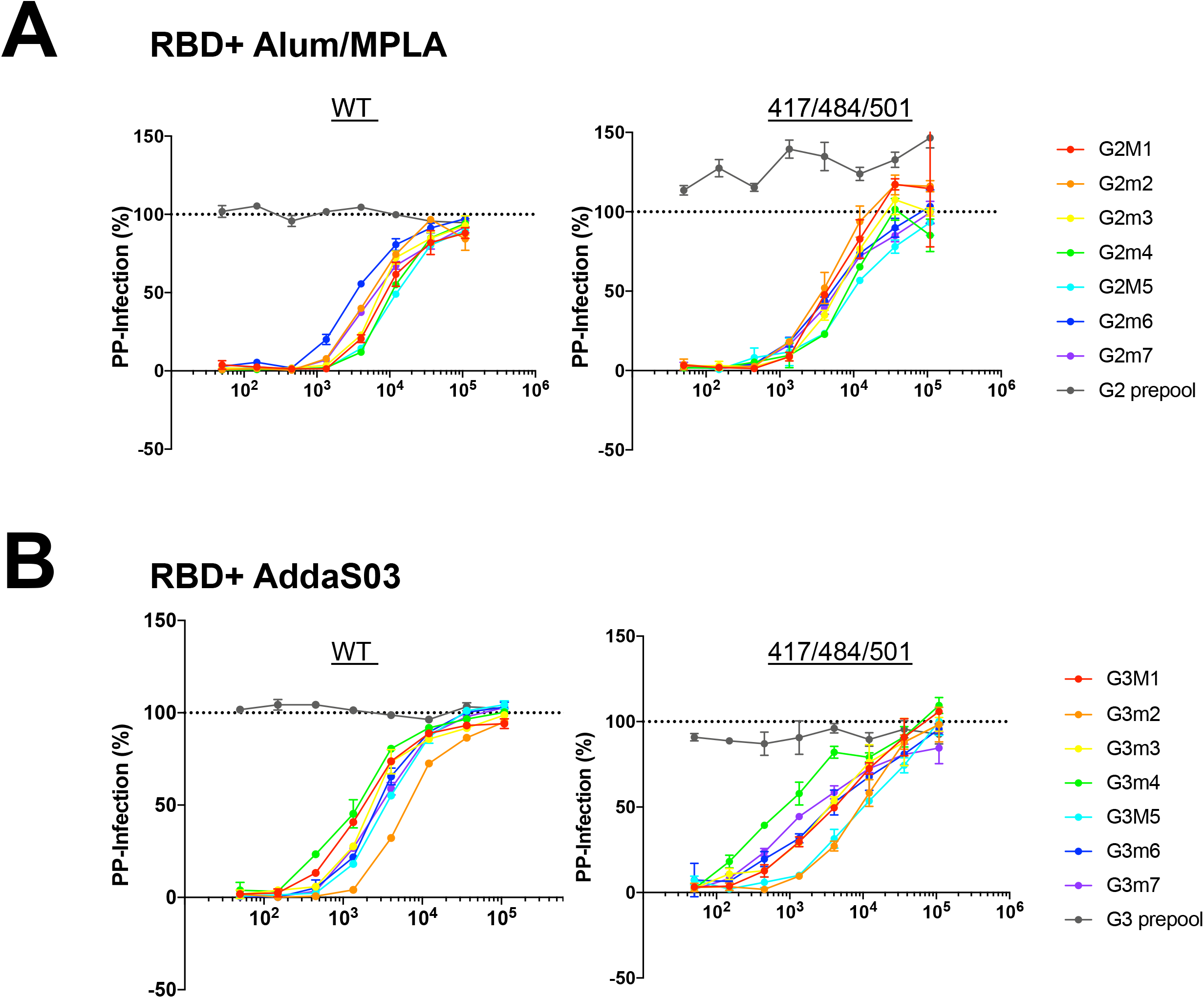
Dose-dependent neutralization activity against SARS-CoV-2pp containing variant mutations N501Y/E484K/K417N. Neutralization activity of mouse sera were tested at 1:50 to 1:109,350 in 3 fold dilution against lentivirus particles pseudotyped with either spike of SARS-CoV-2(WT) or S encoding triple mutations N502Y/E484K/K417N. Pre- (pooled) or Post-vaccination sera were pre-incubated with pseudoparticles (PP) encoding different surface proteins for 1 hour followed by addition to 293 ACE-2 cells. 48 hour post-transduction, entry of PP were quantitated by luciferase activity. Result are done in duplicate normalized to entry of PP in presence of PBS. Representative of two independent experiments is shown.

